# Chaotic dynamics for homeostatic hematopoiesis

**DOI:** 10.1101/2024.08.16.608266

**Authors:** Dongya Jia, Emanuel Salazar-Cavazos, Timothy West, Shen-Huan Liang, Raquel Costa, Maria Clavijo-Salomon, April Huang, Giorgio Trinchieri, Michail Lionakis, Ratnadeep Mukherjee, Grégoire Altan-Bonnet

## Abstract

Homeostatic hematopoiesis is a dynamical process, characterized by large variations (e.g., with coefficients of variation larger than 1) of cell quantities, cell proliferation rates, and extensive correlations/anticorrelations between cell types within the myeloid/lymphoid lineage, and between lineages. All cell types exhibit rare but synchronized bursts of proliferation in the bone marrow, blood and spleen. Through longitudinal study of the blood contents of healthy mice, we found that leukocyte fluctuations are ergodic yet subject to chaotic behaviors characterized by a broad spectrum of characteristic timescales. We then built a minimal mathematical model to capture these dynamical features of hematopoiesis (fluctuations, correlations, and chaos) and explain how the accumulation of B cells (e.g. during lymphoma development) would transition the blood cell dynamics from chaos to oscillations as observed clinically. Finally, we demonstrated the ubiquity of the correlated blood cell fluctuations by comparing mouse cohorts of different genetic backgrounds and ages.

## INTRODUCTION

Classical studies in hematopoiesis have delineated the lineage tree structure underlining the ontogeny and diversity of blood cells. In our textbook understanding, almost all blood-derived leukocytes originate from hematopoietic stem cells (HSC) in the bone marrow, whose differentiation implies lineage split and fate bifurcation resulting in varied cell types. For example, there exists an irreversible split between common lymphoid progenitors (CLP) and common myeloid progenitors (CMP) accounting for the production of natural killer (NK), B and T cells *vs.*, macrophages, dendritic cells and neutrophils^1^. However, this classical understanding has been derived mostly by genetic perturbation (e.g., knocking out “master regulator” transcription factors (TFs), resulting in a loss of entire branches of the hematopoietic tree) or reconstitution (e.g., ablation by irradiation followed by bone marrow reconstitution): both techniques, while tremendously informative, especially for the field of transplantation, may be too perturbative to inform us about hematopoiesis at homeostasis. Recently, novel molecular techniques have unleashed a fresh look at hematopoiesis by allowing the analysis of cell formation in situ without perturbations. ‘Barcoding’ strategies including viral barcodes^2^, transposon insertion site barcoding^3^ and artificial DNA recombination locus^4^ have been developed to tag cells of interest in situ, to track their clonal dynamics, and to assess cell fates. These studies were consistent with the classical model of hematopoiesis as a hierarchical tree of cell differentiation; these provided further insights into the HSC hierarchy by revealing functionally distinct HSC subtypes and long-lived progenitors and refined our understanding of the split between myeloid and lymphoid cells.

The dynamic nature of hematopoiesis remains challenging both conceptually and functionally. There are roughly 330 billion cells turning over every day in a human body, 86% of which are blood cells^5^. Accordingly, hematopoietic homeostasis cannot be considered as a static process, as this would obscure the temporal dynamics of cells and cell number variation associated with such rapid turnover events. This provides the immune system with a highly dynamic population of cells that can fight varied pathogenic onslaughts (viral, bacterial or fungal), with different dynamics, in different tissues. In turn, such consideration challenges the concept of homeostasis: does the hematopoietic system achieve tight regulation of its composition, or does it allow for fluctuations? In that context, previous clinical studies have reported large fluctuations of cell numbers in both patients and healthy volunteers. Spontaneous oscillations of overall white blood cells (WBCs) and neutrophils have been observed in chronic myelogenous leukemia (CML) patients and neutropenic patients. These oscillations were stable for years. The fluctuation of blood neutrophils has also been reported in healthy volunteers^6–8^. Altogether, this suggests that the cell numbers in both physiological and pathological conditions are vibrantly fluctuating with large yet controlled variation. How this clinical observation bears on our understanding of hematopoiesis remains elusive. An intuitive explanation of cell number variation is the stochasticity arising from HSCs due to their small population size^9^. If so, we would expect propagation of noise from HSC to differentiated cell types, which is not the case as we show later.

The tree structure of hematopoiesis offers a hierarchical model of the precursor-progeny relationship during development, but this “genealogical” structure may obscure how “unrelated” cell populations interact with each other (e.g. by competing for space and resources, by synergizing in their generation), missing fundamental regulations within the hematopoietic system. The past two decades have witnessed the advance in our understanding of cell-cell interactions, with the systematic analysis of ligand/receptor interactions^10,11^. However, we are still missing a holistic characterization of cell type correlations during hematopoiesis.

In this study, we revisited fundamental aspects of hematopoiesis at homeostasis in healthy mice. We relied on high-dimensional cytometry (mass cytometry and spectral flow cytometry) to carry out a cross-sectional and longitudinal study of variations in hematopoietic contents in the bone marrow, blood, and spleen of mice. When collecting samples, we made sure to do so at the same set time during the day (10 a.m.) to exclude the effect of circadian rhythm on cell number fluctuation. We applied the Elowitz method^12^ to deconvolve the sources of noise in this system and relied on modified uracil pulsing to quantify cell proliferation. Our experimental measurements ascertain unexpected chaotic fluctuations in leukocyte frequencies at homeostasis, leading us to propose a biochemically realistic simple model of competing leukocytes that accounts for the key features of hematopoiesis revealed in our study (fluctuations, correlation, and chaos), as well as in previous studies in clinical settings (oscillations for leukemia patients).

## RESULTS

### All cell types exhibit large variation of cell frequency in bone marrow, blood, and spleen

To quantify hematopoietic cell frequency we acquired bone marrow, blood, and spleen samples of 39 healthy C57BL/6 (B6) mice (**Fig. 1A**). These inbred mice share identical genomic background and environment (food- and microbiome-wise). We performed mass cytometric (CyTOF) measurements, using a 31-marker panel (**Table S1**) designed to characterize the main hematopoietic cell types at single-cell resolution and assembled a dataset including 6,405,201 cells. To characterize cell types, we applied HAL-x, our custom, scalable, and hierarchical clustering algorithm^13^. A main advantage of HAL-x is its ability to train a clustering model on a subset of cells (e.g. 10^5^), and then to rapidly apply the trained model for clustering analysis of the entire dataset (*>* 10^7^ cells) and/or new datasets. The cells could then be clustered into 29 distinct cell types (**Figs. 1B**−**C**). Overlaid marker expression plots confirm the positions of the identified clusters in UMAP space across all three compartments (**Figs. S1-3**). The most populated cell types are B cells, neutrophils, monocytes, and T cells. (**Fig. 1B**). We found that the bone marrow, blood and spleen contain all the 29 cell types but with various proportions (**Fig. 1C**). For example, the CD93^+^CD19*^−^*B220^+^ cells (B cell precursors, cluster 18) are more populated in bone marrow relative to blood and spleen. In contrast, the CD93*^−^*CD19^+^B220^+^ cells (mature B cells, cluster 28) are more populated in blood and spleen relative to bone marrow. Generally, cell frequency distributions in bone marrow, blood and spleen can be well approximated as log-normal distributions (**Figs. S4-6**). Hand-gating analysis of this CyTOF dataset yielded results that were consistent with our automatic clustering analysis for key cell types (**Fig. S7**).

**Figure 1.**
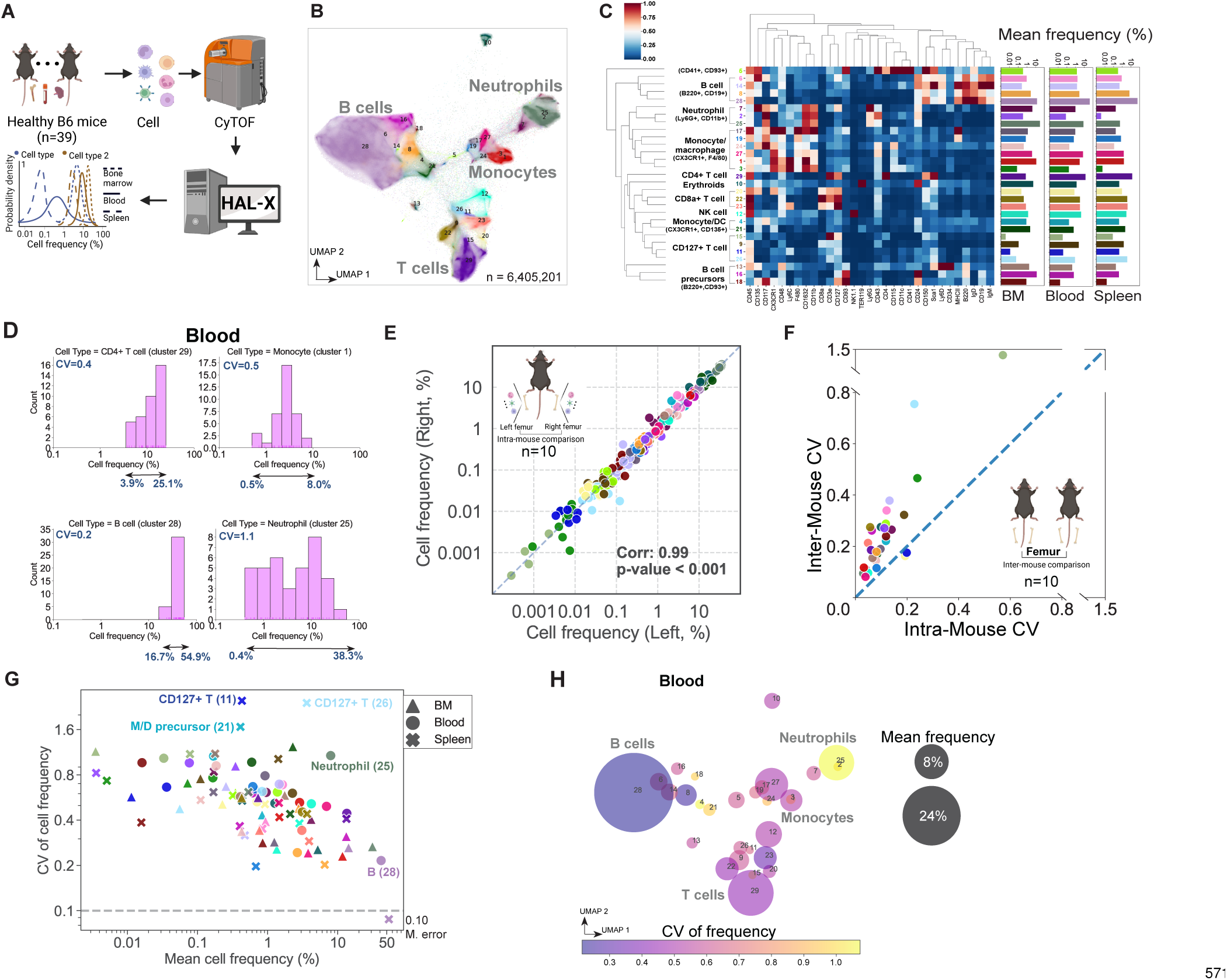
Quantitative analysis of cell number variation in healthy B6 mice. **(A)** Schematic illustration of the pipeline to quantify cell number variation. The bone marrow, blood and spleen samples were collected from 39 healthy B6 mice. As a result, 6,405,201 cells were acquired, followed by CyTOF measurement, HAL-x clustering analysis and cell frequency quantification. **(B)** UMAP visualization showing the majority of the cells being grouped into four main leukocyte types - B cells, T cells, monocytes and neutrophils. We kept a consistent color code for each cell clusters across the rest of this figure. **(C)** Hierarchical clustering result of marker expression showing 29 distinct cell types, where each row represents a cell cluster and each column represents a cell marker. **(D)** Frequency distribution of the CD4^+^ T cells (cluster 29), monocytes (cluster 1), B cells (cluster 28) and neutrophils (cluster 25) in the blood. We present the coefficients of variation (CV) for each cell cluster. **(E)** Individual mouse exhibits close to identical cell frequency for all cell types when comparing left and right femurs. Each dot represents one cell type within one mouse in panels E, F and G. **(F)** The inter-mouse variability is always larger than the intra-mouse variability. **(G)** Large variations in cell frequencies are generic across all cell types in bone marrow, blood and spleen. The dotted line represents the mean intra-mouse CV. **(H)** Visualization of the variations in cell frequencies in the blood. Each blob represents a cell cluster centered according its UMAP coordinates in (B); the blob size is scaled according the mean frequency of the cell type; and the color of the blob maps the CV of the frequency. Large variations in cell number variations are observed in both infrequently–populated cell types (e.g., monocyte/DCs, cluster 4) and highly–populated cell types (e.g., neutrophils, cluster 25).

To quantify the variation in cell frequencies, we calculated the coefficient of variation (CV), defined as the ratio of standard deviation to mean^14^. For example, the frequency of Ly6G^+^CD11b^+^ cells (blood neutrophils, cluster 25) varies from 0.4% to 38.3% across the 39 mice, yielding a CV of 1.1, while the frequency of CD93*^−^*CD19^+^B220^+^ cells (blood B cells, cluster 28) varies from 16.7% to 54.9%, yielding a CV of 0.2 (**Fig. 1D**). In order to rule out that the measured variations in cell frequency were due to measurement and/or data processing errors, we applied the Elowitz method to distinguish intrinsic and extrinsic noise in biological systems^12^: we compared the intra-mouse and the inter-mouse variability by measuring the correlation in cell type frequency between the left and right femurs of a given mouse, and by reporting the variability across mice.

We found that each cell type exhibited nearly identical cell frequency between left and right femurs in each mouse (Pearson correlation: *r_p_*=0.99, *p <*0.001) (**Fig. 1E**). We observed that, even for the very infrequently populated cell types (cluster 15, subset of CD127^+^ T cells), we obtained an excellent match between left (CD127+ T cells, cluster 15, 0.000285%) and right femurs (CD127+ T cells, cluster 15, 0.000279%) (left bottom of (**Fig. 1E**). Such observation demonstrates that our protocol to estimate the fluctuations in hematopoietic frequencies (including cell isolation, staining, cytometric characterization and computational analysis) was practically noise-free. We found the inter-mouse variability of all cell types to be always larger than the intra-mouse variability (except for clusters 20, subset of CD8*α*^+^ T cells and cluster 11, subset of CD127^+^ T cells which have comparable intra-mouse and inter-mouse variability) (**Fig. 1F**). Hence, to paraphrase the framework introduced by Elowitz *et al.*^12^, variations in the abundance of leukocytes are driven by extrinsic factors rather than intrinsic noise. Additionally, these left/right femur correlations for all hematopoietic cell types hint at a global mode of regulation. Altogether, these dual measurements within individual mice established that the large variations of cell frequency from mouse to mouse are not due to measurement/processing error, but, rather and more fundamentally, are of biological origin.

In bone marrow, blood and spleen, we found that all cell types exhibit large variation of frequency, exemplified by the blood Ly6G^+^CD11b^+^ cells (neutrophils, cluster 25) and the spleen CD127^+^ T cells (clusters 11 and 26) (**Fig. 1G**). The inter-mouse variability of cell frequency is characterized by CVs varying between 0.1 to 2.5 with the median being 0.5 in bone marrow and spleen, and 0.6 in blood, indicating a typically more than 10-fold variation (**Fig. 1D**). Notably, the cell types that displayed large variations include both infrequently populated cell types, such as blood monocyte/DCs (cluster 4 with a mean of 0.17% and a CV of of 1.1) and highly populated cell types, such as blood neutrophils (cluster 25 with a mean of 7.93% and a CV of 1.1) (**Fig. 1H**). Similar large variations in cell frequency can also be obtained by hand-gating the CyTOF data and defining cell types in a classical manner (**Figs. S8A-C**). Notably, there is no hierarchy in frequency variations derived from variations in HSC frequencies, as precursor cell types can have higher or lower CVs relative to their progenies **(Fig. S8D)**.

We then theoretically explored whether the large variations in cell frequency were a consequence of intrinsic (stochastic nature of cell dynamics) and/or extrinsic cellular noise (fluctuation of cell kinetics etc.). First, we considered the potential impact of intrinsic noise *i.e.* how stochasticity in molecular/cellular processes may result in variation. To have a solid understanding of how intrinsic noise alone may affect cell number variation in the hematopoiesis, we used a mathematical model focusing on the dynamics of one cell type by considering three fundamental processes – production Ω, proliferation *λ*, and disappearance *d* - through death or further differentiation (**SI Section 1**). We derived the master equation representing the time evolution of cell numbers and solved the master equation analytically by the generating function technique^15^. We derived analytical formulae for the mean cell number *< n >*= Ω*/*(*d* − *λ*) and the variation 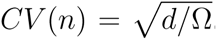. Notably, this implies that the cell proliferation rate *λ* does not impact the variation in cell numbers: changing *λ* rescales the standard deviation and the mean of cell number to the same extent and thus doesn’t change their ratio – CV. We then explored the biological requirements for the rates (Ω,*⋋*,*d*) to match the experimentally measured cell statistics. We focused on the bone marrow neutrophil (cluster 7) as a striking example of the limitations of the intrinsic-noise-only model: mouse neutrophils have a half-life of around 8 hours^16^, and therefore the decay rate is ln2/8=0.09 *hour^−^*^1^. The CV of bone marrow neutrophil (cluster 7) being estimated to be 0.4, one can then estimate Ω = *d/CV* (*n*)^2^ = 0.6 *hour^−^*^1^. The total number of mouse bone marrow cells are 10^8^ ^17^ and the mean frequency of cluster 7 is 4%. Therefore, *d* − *⋋* = Ω*/ < n >*= 1.5 ∗ 10*^−^*^7^. To fulfill the large mean number *< n >* and large variation *CV* (*n*) of bone marrow neutrophils, mice would have to maintain the disappearance *d* and proliferation rate *⋋* at an extremely high degree of precision -*λ* would need to be equal to *d* within (*d* − *λ*)*/d* = 2.5 ∗ 10*^−^*^7^. This is biologically incongruous and incompatible with our measurements, as demonstrated in the following section of quantifying cell proliferation rates.

### All cell types exhibit large variation of proliferation rates

Systematically measuring the fluctuation of cell kinetics (differentiation, proliferation or death) in hematopoiesis would imply a complex longitudinal kinetic study beyond the experimentally feasible at present time. However, we could estimate an upper-bound for the variation of cell proliferation rates for all cell types under study. We used a 5-iodo-2’-deoxyuridine (IdU)-pulsing protocol by performing intraperitoneal injection of IdU with subsequent analysis by mass spectrometry (**Fig. 2A**). Note that such injection does not induce any detectable inflammation and does not induce perturbations of the hematopoietic homeostasis^18^. As different cell types in mouse exhibit comparable duration of *S* phase in cell cycle^19^ - in other words, once cells start replicating their DNA, S-phase proceeds like clockwork - then we could estimate cell proliferation rates by measuring IdU positive (IdU^+^) fractions of that cell type (**SI Section 2**). To rule out the possibility that the mouse-to-mouse variations is due to measurement/data processing errors, we again applied the Elowitz method^12^ to quantify the intra-mouse variability of cell proliferate rates. We showed that the same cell type exhibited almost identical (*r_p_*= 0.97*, p <* 0.001) proliferation rates between left and right femurs within a given mouse (**Fig. 2B**). Each cell type exhibited much larger inter-mouse CV than intra-mouse CV of cell proliferation rates (**Fig. 2C**). Altogether, our measurements confirmed that the large variation of cell proliferation rates is of biological origin, and not a result of experimental artefacts.

**Figure 2.**
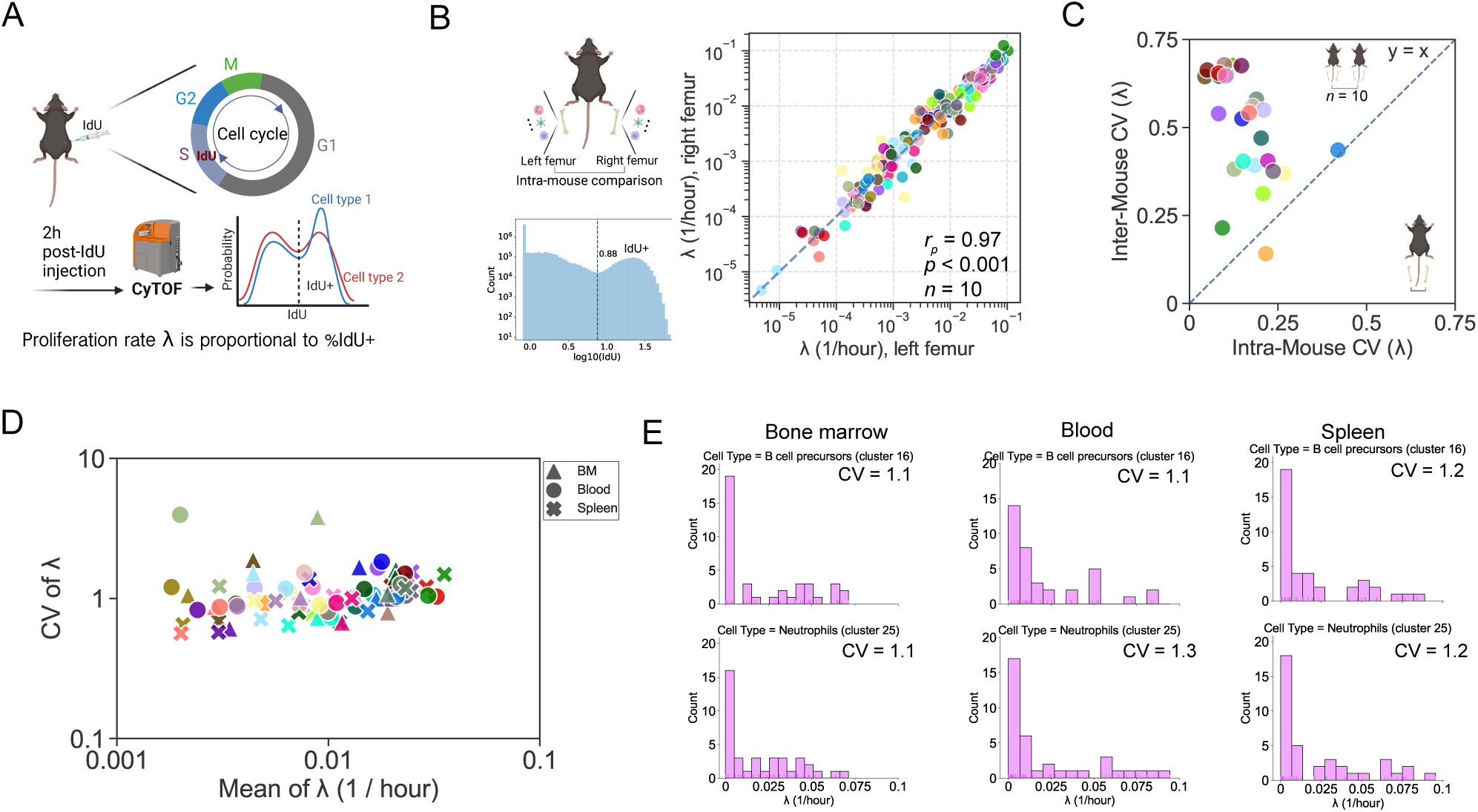
Quantifying the variation of cell proliferation rates in healthy B6 mice. **(A)** Schematic illustration of the IdU-pulse protocol to estimate cell proliferation rate (*λ*). **(B)** Cell proliferation rates exhibit close to identical correlation between left and right femurs in individual mice. Each dot represents a cell type in panels B-D. **(C)** The inter-mouse variability of cell proliferation rates is significantly larger than the intra-mouse variability. **(D)** Cell proliferation rates exhibit large variations with coefficients of variation around one, across all cell types in bone marrow, blood and spleen. **(E)** Note the ”long-tails” distribution of the cell proliferation rates of B cell precursors (cluster 16) and neutrophils (cluster 25). These two cell clusters have comparably large variations across bone marrow, blood and spleen.

We found that all 29 cell types in all three compartments (bone marrow, blood and spleen) exhibited large variation of proliferation rates characterized by CVs with median being 1 (**Fig. 2D**). Interestingly, the distribution of the proliferation rates of each cell type had ‘long tails’, demonstrating that there always exists a small proportion of mice undergoing burst of cell proliferation (**Fig. 2E**). For example, the CD93^+^B220^+^ cells (subset of B cell precursors, cluster 16) and the Ly6G^+^CD11b^+^ cells (subset of neutrophils, cluster 25) exhibited comparable CVs of proliferation rates in bone marrow, blood and spleen (**Fig. 2E**). Consistent large variations in cell proliferation rates were obtained by hand-gating the CyTOF data and defining cell types in a classical manner (**Figs. S9A-C**). Interestingly, multipotent progenitors (MPPs) displayed the highest proliferation rate on average (**Fig. S9D**) and HSCs had the highest variability of cell proliferation rate (**Fig. S9E**). Strikingly, in 3 out of the 39 mice, more than one third of their blood neutrophils (cluster 2) were found to be IdU^+^ i.e. to be in the midst of DNA replication and cell proliferation (**Fig. S10**). We further connected the CV of cell proliferation rate with the CV of cell frequency and found that there is a small yet significant positive correlation, confirming that fluctuating cell proliferating rates may contribute to the uncovered variation in cell frequencies (**Fig. S11**).

In summary, individual cell types exhibited large variations of both frequency and proliferation rate, which appears incompatible with our classical understanding of homeostatic hematopoiesis. To be explicit, rather than fixed rates of differentiation/proliferation/death, mice seem to experience large fluctuations of rates (at least for proliferation), complementing our observation that intrinsic stochastic noise in cellular dynamics might be insufficient to account for the large observed variations in leukocyte frequencies. We then proceeded to further explore the ‘structure’ underlying the variations by connecting cell types.

### Extensive correlations/anti-correlations exist between cell types within myeloid lymphoid lineage and between lineages

We next investigated whether the large fluctuations in hematopoietic cell frequencies were consistent with the “tree” view of cell generation in our textbook understanding. We reasoned that there should be significant correlations based on the differentiation hierarchy (**Fig. S12A**). We revisited the classical deterministic mathematical model of hematopoiesis introduced by Bariel *et al.*^20^: in this study, the kinetic parameters (proliferation, differentiation and death) of 16 cell types were carefully quantified using fate-mapping of HSC and BrdU accumulation. These measurements and model demonstrated that HSCs mainly account for recovery of hematopoiesis under stressed conditions, while MPPs mainly sustain native hematopoiesis at homeostasis. We carried out stochastic simulations to evaluate the time evolution of population size of the 16 cell types and further characterized their correlations. Starting with 1000 HSCs, we performed the simulation representing a 120-day dynamics by applying Gillespie algorithm to the mathematical model and kinetic rates presented in^20^. We then calculated the Spearman correlation (*r_s_*) among cell types, and found that such hematopoietic tree structure generates a distinct modular correlation pattern: (1) the strong positive correlations are mainly existing separately within the myeloid lineage or within the lymphoid lineage; (2) the strong negative correlations are mainly existing between progenitors of erythroid cells/basophils and the lymphoid lineage; (3) the myeloid and lymphoid lineages exhibit much less pronounced correlations (**Fig. S12B**). To test the model-predicted correlations, we re-analyzed our CyTOF dataset using manual gating of the cell types in the lineage tree (**Fig. S12A**) and found that significant positive and negative cell frequency correlations exist *within and between* myeloid and lymphoid lineages (**Fig. S12C**). There are several modules containing cells that are strongly positively correlated, including (pro B, pre B, imm B and B cells), (HSC-LT, HSC-ST, MEP), (CLP, CMP, BLP), (DN T, DP T, CD4 T and CD8 T) and (MPP2, MPP3, MPP4, MDP, cMoP, GMP, pre-pro B) (**Figs. S8F, 12C**). Hence, our measurements appear not to comply with a purely–stochastic origin for the fluctuations in hematopoietic cell frequencies.

To estimate the cell-cell correlations with improved resolution, we evaluated the 29 cell types identified by clustering in Fig. 1B. We found extensive significant positive or negative correlations of cell frequencies in the bone marrow (**Fig. 3A**), blood (**Fig. S13A**) and spleen (**Fig. S13C**). The positive or negative correlations of cell frequencies exist within lymphoid or myeloid lineages. In our datasets acquired with 39 mice, we performed bootstrapping and confirmed the correlation results to be statistically robust (**Fig. 3A**). For example, in the bone marrow, the frequency of the CD117^+^CD93^+^CD19*^−^*B220^+^ cells (B cell precursors, cluster 18) is positively correlated with the frequency of the CD117*^−^*CD93*^−^*CD19^+^B220^+^ cells (mature B cells, clusters 8) (**Fig. 3A**), consistent with their differentiation hierarchies. On the other hand, the two subtypes of monocytes − the Ly6C*^−^*MHCII^+^ cells (cluster 19) and the Ly6C^+^MHCII*^−^* cells (cluster 3) are negatively correlated (**Fig. 3B**). More substantially, we uncovered strong positive and negative correlations between lymphoid and myeloid lineages. For example, there is a strong positive frequency correlation between CMP and CLP (**Fig. S8F**). Other examples include CD117*^−^*CD93*^−^*CD19^+^B220^+^ cells (mature B cells, cluster 8) and the CX3CR1^+^CD135^+^ cells (macrophage/DC precursors, cluster 21) (**Figs. 3A, S14**). The strongest negative correlation in the bone marrow is between the CD135*^−^*Ly6G^+^CD11b^+^ cells (differentiated neutrophils, cluster 25) and the CD117*^−^*CD93*^−^*CD19^+^B220^+^ cells (differentiated B cells, cluster 8) (**Figs. 3A, C**). The pronounced positive (*r_s_ >*= 0.7) and negative (*r_s_ <*= −0.7) correlations are shown in (**Fig. 3D**). These cell-cell correlation patterns are consistent across tissues. For example, the mature B cells (cluster 8) and the mature neutrophils (cluster 25) exhibited a negative correlation in both bone marrow (**Fig. 3A**) and blood (**Fig. S13A**). Monocytes (cluster 19) and B cell precursors (cluster 13) exhibit a positive correlation in both bone marrow (**Fig. 3A**) and blood (**Fig. S13A**). Moreover, we found cell types, *e.g.*, neutrophils (cluster 25), exhibiting significant positive correlation across bone marrow, blood and spleen (**Fig. 3E**), indicating the inference potential of bone marrow neutrophil levels by blood neutrophils and suggesting the cell-cell correlation is not just a consequence of cell trafficking and organ compartmentalization.

**Figure 3.**
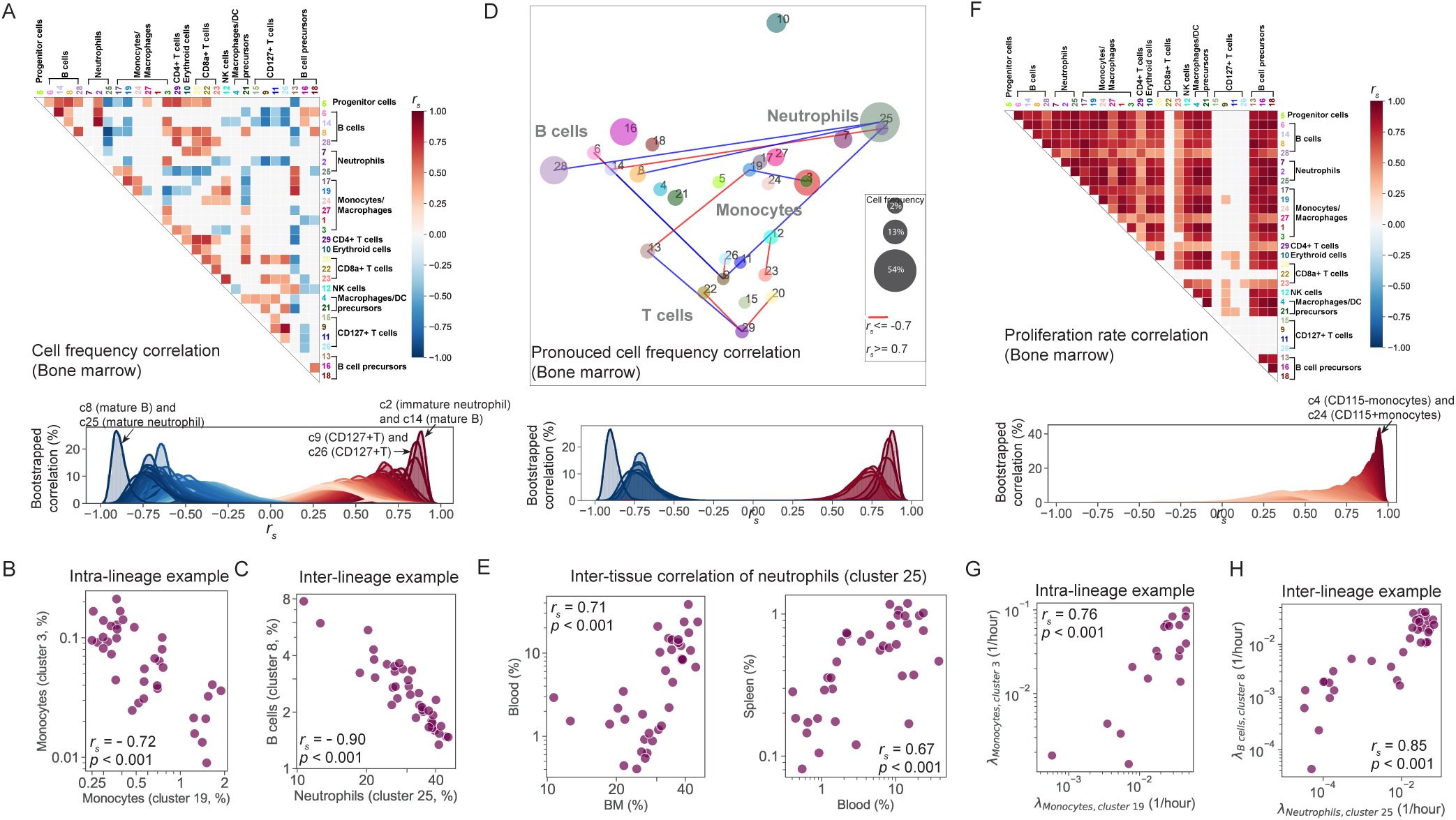
Extensive positive/negative correlations of cell type frequencies exist within and between myeloid and lymphoid lineages in the bone marrow. **(A)** Top: correlation matrix of the fluctuating cell frequencies showing the significant positive (in red) and negative (in blue) Spearman correlation coefficients between all cell types in the bone marrow (similar graph for blood and spleen can be found in the Supplementary Materials **Fig. S13**). Bottom: For each of the significantly correlated cell types, we performed a bootstrapping analysis with 10,000 iterations and obtained the distribution of the correlation coefficients. **(B)** Two subsets of monocytes (clusters 3 and 19) exhibit significantly negative correlation. Each dot in panels B–E and G–H represents cell frequencies within one mouse.**(C)** B cells (cluster 8) and neutrophils (cluster 25) exhibit significantly negative correlation. **(D)** Visualization of leukocyte fluctuations for the strongest positive (n = 7) and negative (n = 7) correlated pairs, as defined by the absolute value of the correlation coefficient being larger than 0.7 (with the centroids of each cell cluster centered on UMAP coordinates reported in Fig. 1B. **(E)** Neutrophil frequencies exhibit significant positive correlation across bone marrow, blood and spleen. **(F)** Top: The correlation matrix showing the significant Spearman correlation coefficients of cell proliferation rates of all cell types in the bone marrow. Strikingly, there are only positive correlations depicted in red color. Bottom: For each of the significant correlated cell proliferation rates, we performed bootstrapping analysis with 10,000 iterations. The distribution of the correlation coefficients is shown in the bottom panel. **(G)** The proliferation rates of the two subsets of monocytes (cluster 3 and 19) are significantly positively correlated. **(H)** The proliferation rates of B cells (cluster 8) and neutrophils (cluster 25) are significantly positively correlated.

Altogether, these results suggest that our textbook understanding of hematopoiesis^20,21^ is not sufficient to explain the extensive cell-cell correlations in our measurements. We conjectured that such discrepancy may stem from extrinsic factors altering the dynamics rates along the lineage tree of differentiation, accounting for the negative correlations within the same lineage and positive correlations across lineages. Given the extensive frequency correlations between cell types, we then explored whether cell frequency correlations could be explained by the strongly– correlated fluctuations in the cell proliferation rates.

### The hematopoietic system undergoes synchronized bursts of proliferation

Among cell proliferation rates, we found that, if there is a significant correlation, it is only positive (**Fig. 3F**). Even between the cell types that exhibit anticorrelated frequencies, e.g., two subtypes of monocytes (clusters 3 and 19) (**Fig. 3B**) and B cells and neutrophils (clusters 8 and 25) (**Fig. 3C**), their proliferation rates are positively correlated (**Figs. 3G-H**). We also observed this global positive correlation pattern of cell proliferation rates in the blood (**Fig. S13B**) and spleen (**Fig. S13D**). Note that our measurements were time-synchronized at 10am, within a cohort of female mice sharing living quarters: this ruled out diurnal and menstrual cycles, as well as microbial cues, as potential sources of proliferation burst. The global positive correlation pattern of cell proliferation rates suggests that the hematopoietic system undergoes rare but synchronized burst of proliferation (with most likely concomitant burst of cell death to maintain homeostasis). We further confirmed that the cell proliferation rates do fluctuate in a synchronized manner across cell types, using an orthogonal approach of intracellular staining of the cell proliferation marker Ki-67 (**Fig. S15A**).We acquired one to two drops of blood from 24 healthy B6 mice, and immediately processed the samples and performed spectral flow cytometry using a panel of 18 surface markers, followed by permeabilization and intracellular staining of Ki-67. We found high consistency in characterizating the variation of cell proliferation rates between the measurement by IdU incorporation and the measurement by Ki-67 staining. We found that the distribution of %Ki-67^+^ exhibit ’long-tails’ consistently with our IdU measurements (**Figs. S15B-C**). We showed that the variation of cell proliferation rates across mice identified by Ki-67 staining (median (CV) = 1.26) (**Fig. S15D**) is comparable with the analysis result by IdU Pulse (median (CV) = 1.05, **(Fig. 2)**. We showed that there is only positive correlation between %Ki-67^+^ of different cell types (**Fig. S15E**), confirming the synchronized burst of hematopoietic cell proliferation.

We conjectured that these synchronized bursts of cell proliferation may stem from the pleiotropic effects of cytokines. It has been speculated that while the production of these soluble ligands is probably cell-type-specific, the binding of ligands across blood cells is promiscuous ^22^. Additional details have been revealed in two recent studies: Cui et al. identified more than 66 cytokines that drive cellular polarization across immune cell types^11^. Orcutt-Jahns et al. reported that IL-10 up-regulates pSTAT3 and IL-4 up-regulates pSTAT6 across 23 different cell types (including T cell, B cell, myeloid cells etc.) in human peripheral blood mononuclear cells^23^. These studies highlight how small changes in the cytokine milieu may perturb a large swath of leukocytes (with higher/level of cytokine receptors and/or proliferation/disappearance), with potentially knock–on effects across the whole hematopoietic tree^24^.

### A longitudinal study demonstrates the ergodicity of the hematopoietic system and reveals the potential chaotic nature of its homeostasis

To further our understanding of the uncovered hematopoietic fluctuations at homeostasis, we performed a longitudinal study to reveal the blood dynamics in homeostasis. We acquired, every two or three days, one to two drops of blood from 10 healthy B6 mice at a fixed time (noon), followed by immediate processing and spectral cytometry (CyTEK) analysis for 15 cell surface markers covering cell types including neutrophils (Ly6G^+^), B cells (CD19^+^), T cells (CD3^+^) and monocytes (CD11b^+^) (**Fig. 4A, Table S2**). In total, we acquired the blood profiling results at 12 different days over a one-month period. Certain cell types being fragile and prone to apoptosis, *e.g.*, neutrophils, we optimized our protocol (including red blood cell lysis, antibody staining and cell fixation) to minimize the time from blood collection to cell fixation under one hour and a half: such optimized protocol was critical to reduce cell loss and to circumvent limitations in previous analysis of the hematopoietic system (**Fig. S16**).

**Figure 4.**
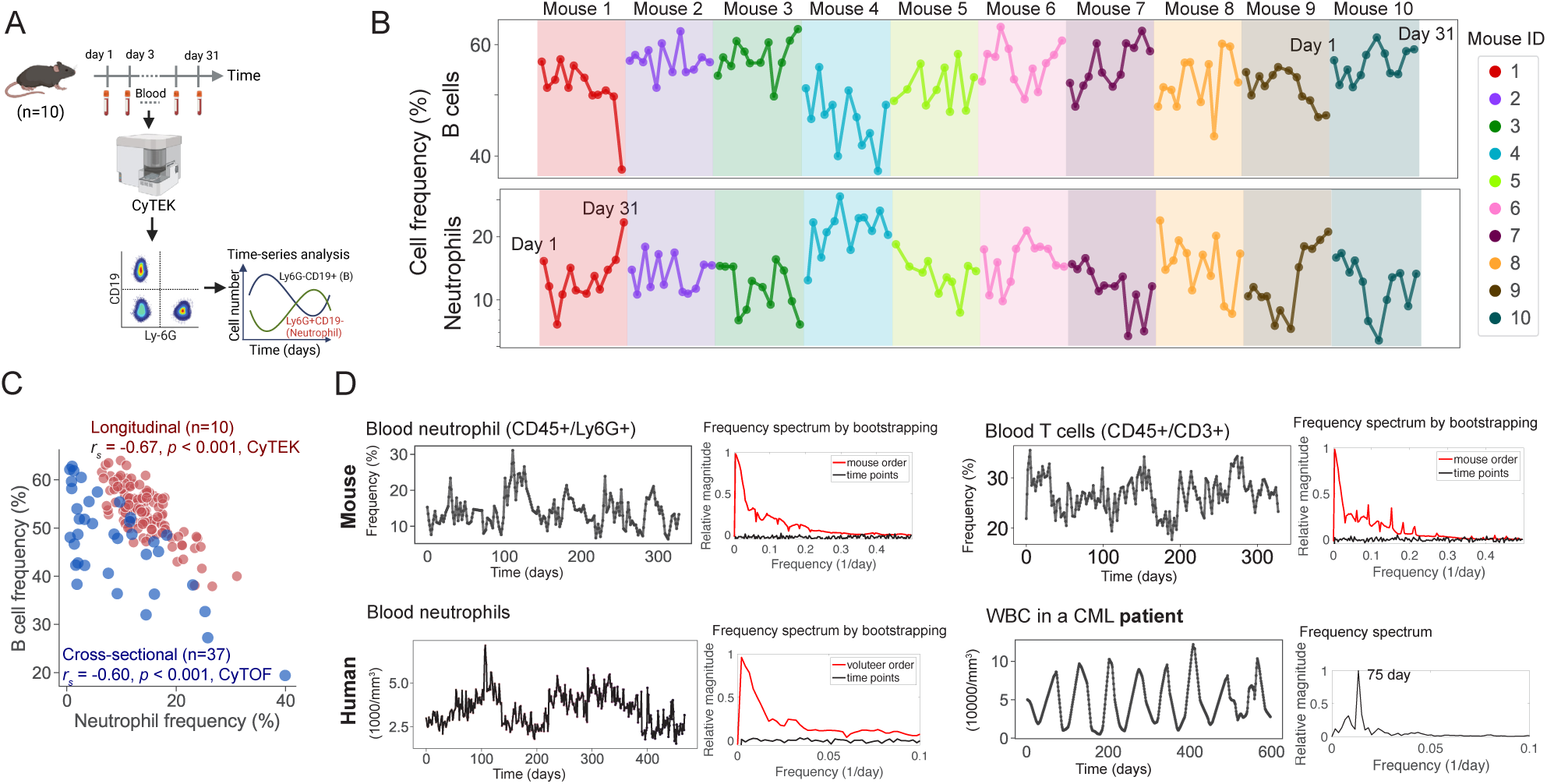
A one-month longitudinal study of mouse blood profiling. **(A)** Schematic illustration of the longitudinal study: one to two drops of blood were acquired every two to three days from 10 B6 mice, followed by immediate processing/measurement by spectral flow cytometry and hand-gating for blood neutrophils (CD45^+^Ly6G^+^) and B cells (CD45^+^CD19^+^). **(B)** Concatenated time-series of mouse neutrophils and B cells in the blood of each of the 10 mice over a month. Top: fluctuation of B cell frequency; Bottom: fluctuation of neutrophil frequency. **(C)** B cells and neutrophils in mouse blood exhibit comparable negative correlation in the present longitudinal relative to the previous transversal analysis of healthy mice (Fig. 3). **(D)** Fourier spectrum analysis of the fluctuations in hematopoietic cell components reveal a continuum of frequencies in the blood of both healthy mice (top) and healthy human (bottom left), while the fluctuations in white blood count (WBC) for a chronic myeloid leukemia (CML) patient display a single-frequency/peaked Fourier spectrum. In each subplot, the left shows the time-series of cell frequency/concentration, and the right shows the frequency spectrum of the time-series.

We first checked that there is no correlation between the number of CD45^+^ cells and the frequency of neutrophils or the frequency of B cells (**Fig. S17**), demonstrating that the cell frequency is not affected by the total number of CD45^+^ cells analyzed, *i.e.* the volumes of collected blood. We then checked the range of neutrophils and B cells per mouse and all ten mice exhibit comparable variation of cell frequencies (**Figs. S18A-C**). We then checked the range of neutrophils and B cells per time point and found no drift over time, indicating the processing of acquiring one to two drops blood from the mice every two to three days over one month did not alter hematopoietic homeostasis (**Fig. S19A-B**). Yet, we found that within each of the ten mice, the overall B cell frequency and neutrophil frequencies were fluctuating and negatively correlated (**Figs. 4B, S18D**). We confirmed that the negative correlation between B cells and neutrophils could be observed at individual time points **(Fig. S19C)**. The correlation coefficient calculated by the longitudinal study of 10 mice (*r_s_* = −0.67) is comparable to that calculated by the cross-sectional study using 39 mice (*r_s_* = −0.60) (**Fig. 4C**), which itself validates the concept that hematopoiesis is an ergodic process, *i.e.*, the ensemble average and the time average for the hematopoietic population frequencies are identical.

Motivated by the apparent noisiness of the time-series of leukocyte frequencies (**Fig. 4B**), we proceeded to evaluate whether the hematopoietic system should be modeled as a multiple-frequency oscillator or possibly a chaotic system. We computed the Lyapunov exponents and, more specifically, the maximal Lyapunov exponent (MLE) for the hematopoietic dynamics: indeed, if MLE*>*0, the system would display high sensitivity to initial conditions, a hallmark of chaotic dynamics^25^. We assembled an overall continuous time-series of neutrophils and T cells by concatenating the individual time-series of the ten mice and using linear imputation (**Figs. S20-21**). We bootstrapped our analysis of the time series by shuffling the order of concatenation across mice: this demonstrated that our concatenation of time-series did not introduce any bias. For each time-series, we calculated the spectrum of Lyapunov exponents using Eckmann’s algorithm^26^ and performed a Fourier transformation to generate the frequency spectrum. We found that the MLE of the neutrophil time-series is 0.13 ± 0.02 (*mean* ± *std* of 1000 bootstrapping tests) and the Fourier spectrum exhibited continuous frequencies especially for the range of frequency smaller than 0.1/day, *i.e.*, the period larger than 10 days (**Fig. 4D**). We checked that randomization of the time series across time points flattened the power spectrum. Additionally, the time series of blood T cells were characterized by MLE = 0.14 ± 0.03 (*mean* ± *std* of 1000 bootstrapping tests) and with a continuous Fouried spectrum (**Fig. 4D**). Thus, we concluded that the temporal dynamics of blood neutrophils and T cell fluctuations were associated with positive MLEs and continuous Fourier spectra, leading us to conclude that the potential chaotic nature of hematopoiesis in healthy mice.

We performed a similar dynamic analysis for the blood cells in humans. We reconstructed the time-series of neutrophils from eight healthy volunteers from datasets reported in the literature^6^ (**Figs. 4D, S22**) and performed bootstrapping analysis to identify the robust MLE and Fourier spectrum of frequencies. The neutrophil fluctuations in the healthy volunteers exhibited chaotic pattern characterized by a positive MLE (0.16 ± 0.01) and continuous frequencies on Fourier spectrum (**Fig. 4D**), similar to our results in mice. In contrast, the blood cells in patients often exhibited oscillatory fluctuations with a well–defined frequency. For example, the white blood count (WBC) fluctuation in a chronic myeloid leukemia (CML) patient^27^ exhibited very steady oscillations characterized by a period of 75 days (**Fig. 4D**). Note that, during collection of this WBC time-series, no treatment was applied^8^. The loss of frequency of blood cell fluctuation has been observed in the fluctuation of other cell types including platelets and reticulocytes in CML patients (**Fig. S23**) and the fluctuation of blood neutrophils in neutropenic patients (**Fig. S24**) but with very varied and yet unaccounted–for frequencies. All together, our experimental results and our reprocessing of the literature demonstrate that chaos (*i.e.* a sensitivity to initial conditions and deterministic fluctuations) is probably a general pattern for blood cell dynamics in healthy condition in both mouse and human.

### A mathematical model explains the chaotic behavior of anti-correlated cell frequencies with synchronized cell proliferation

Here we illustrate the deterministic yet potentially chaotic nature of hematopoiesis by introducing a mathematical model of the dynamics of leukocyte populations. The phenomenological model focuses on two interacting cell types (e.g., to mimic neutrophil and B cell) and one common cytokine that regulate the proliferation of both cell types (**Fig. 5A**). For each cell type, we considered three fundamental processes, i.e.,(1) cell production from a precursor population, (2) cell decay including natural decay or decay due to cell-cell direct/indirect interactions, and (3) cell proliferation regulated by the cytokine (**Fig. 5A**). The values of parameters are chosen based on our previous experimental work on quantifying cytokine dynamics and cell proliferation^24^, to make sure the cell numbers are within biological ranges. Here we focused on the possible biologically-realistic minimal structure of cell-cell interactions that recapitulates the key features of hematopoietic system. A more detailed explanation of the model formulation and its parameterization can be found in **SI section 3**.

**Figure 5.**
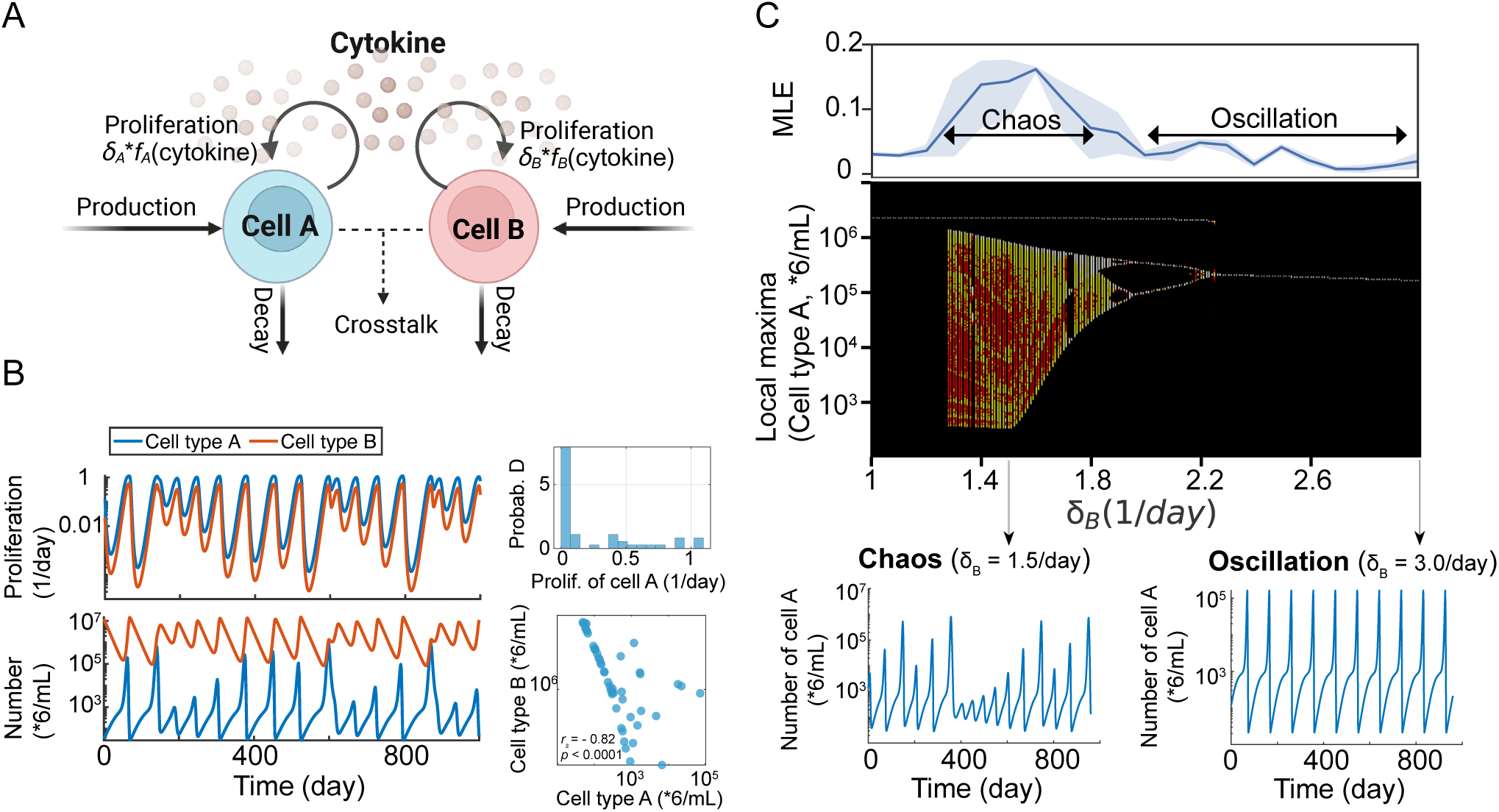
A mathematical model explains the chaotic dynamics characterized by anti-correlated cell frequencies with synchronized cell proliferation. **(A)** Schematic illustration of the structure of the model. The model comprises three components: two cell types (A and B) (considering their production, proliferation, and decay) and one cytokine, which functions as a common resource regulating proliferation of both cell types. **(B)** The two cell types exhibit synchronized proliferation (*top left* ). The distribution of the proliferation rates of cell type A exhibit the ’long tail’ as observed in experiment (*top right* ). The temporal dynamics of quantities of cell types A and B (*bottom left* ). The two cell type exhibit anti-correlated quantities (an example by sampling 50 data points from the time-series (*bottom right* )). **(C)** The dynamic behaviors of the two cell types, exhibiting chaotic pattern. The bifurcation diagram showing the local maximum of the time-series of cell type A as a function of the basal proliferation rate of cell type B *δ_B_* (*middle*). For each *δ_B_*, 100 initial conditions were used for simulation and for each initial condition, the local maxima of cell type A quantity were collected and reported. The maximum Lyapunov exponents (MLEs) of the time-series of cell type A are shown on top of the bifurcation to characterize the chaotic regime (*top*). Examples of increasing *δ_B_*changing the system from chaotic to oscillatory behaviors (*bottom*).

We first explored whether the model recapitulated the experimentally-identified features of cell dynamics. The model generated large CVs in cell frequency and proliferation rate, within the range of our experimental results (**Figs. 1D, 2E, S26A**). The model also recapitulated, by design, the negative correlation between two cell types while their proliferation rates being positively correlated (**Figs. 5B, S26B-C**): this is consistent with our results on neutrophils and B cells *etc.* We then proceeded to learn from the model about possible dynamical features due to such cell-cell interaction structure. Strikingly, our deterministic model with three variables (two cell types and one cytokine) coupled via realistic biological interactions is sufficient to generate chaotic dynamics (**Fig. 5C**): in a nutshell, the competition for a shared cytokine, and its global impact on cell proliferation for two competing cell types can be enough to destabilize an oscillatory system of otherwise cross-inhibiting cells, this results in a strange attractor with a complex power spectrum (**Fig. S26C**) in the dynamics, that is the hallmark of chaotic dynamics. We want to stress that this minimal yet realistic model does generate large fluctuations without invoking stochastic processes, thanks to the chaotic nature of its dynamics.

As abnormal B cell accumulation is typical of leukemia patients, we evaluated how the cell proliferation rate can affect the dynamics in our model. We found that the basal proliferation rate of cell type B (*δ_B_*) can be a critical factor in steering the dynamics. Too fast or too slow basal proliferation rate for B cells can max out (leukemia) or minimize (leukopenia) B cell accumulation, making our three-variable dynamic model collapse to a 2-dimensional problem: such dimensional collapse is enough to drive the system out of the chaotic region and result in regular oscillations (**Figs. 5C**). While such transition between oscillations and chaos is well understood at the mathematical and physiological levels^25,27^, it is, to our knowledge, a new realization in the context of a realistically structured and parametrized model of hematopoiesis. This model prediction explains the clinical observations on the chaotic behavior of neutrophil fluctuations in healthy volunteers and oscillations of WBC, platelets, and reticulocytes in CML patients^8,28^. In particular, it predicts that perturbed hematopoiesis can result in oscillations of very diverse frequencies. Another key parameter predicted by the model is the natural decay rate of cell type B (*γ_B_*) (**Fig. S27**). As delineated by the model, increasing *γ_B_* can lead to the accumulation of cell type A and change the behavior of the system from chaos to oscillations. At the more fundamental level, such a dimensional collapse of the strange attractor of hematopoiesis would be sufficient to explain the transition from chaotic dynamics to oscillatory dynamics, as observed in the clinic ^7,8,28,29^.

### Universal hematopoietic fluctuation and the robust neutrophil-B cell anti-correlation upon treatment

We speculated that the fluctuation and correlation of the hematopoietic components is a general phenomenon. To evaluate how the variation and correlation of cell frequencies persist across different strains of mice, we performed blood profiling analysis of 44 B6 mice including 30 young ones (4-6 weeks old) and 14 aged ones (6.5-months old), together with 30 young (4-6 week old) BALB/c mice, two of the most commonly used inbred strains of laboratory mice. By acquiring one to two drops of blood from each mouse, followed by immediate processing and spectral cytometry analysis for 29 cell surface markers (**Table S3**) covering main leukocyte types, we assembled a dataset of 1,895,969 cells (**Fig. 6A**). The batch effect between B6 and BALB/c samples was minimized as we processed all blood samples side by side (**Fig. S28**). We applied HAL-x^13^ to classify the cells into 19 distinct clusters which belong to four main cell groups - B cells, T cells, neutrophils and monocytes. Both B6 and BALB/c contain all cell types (**Fig. S28** and **Fig. 6B**). The mean frequency of each cell type exhibit a significantly–positive correlation between B6 (young, n = 30) and BALB/c (young, n = 30) mice, with the correlation of highly populated cell types being more pronounced (**Fig. 6B**, bottom left). Notably, the variation of cell frequencies is also significantly correlated between B6 and BALB/c mice (**Fig. 6B**, bottom right), suggesting the variation of cell frequency to be a robust feature that is consistent across different mouse stains.

**Figure 6.**
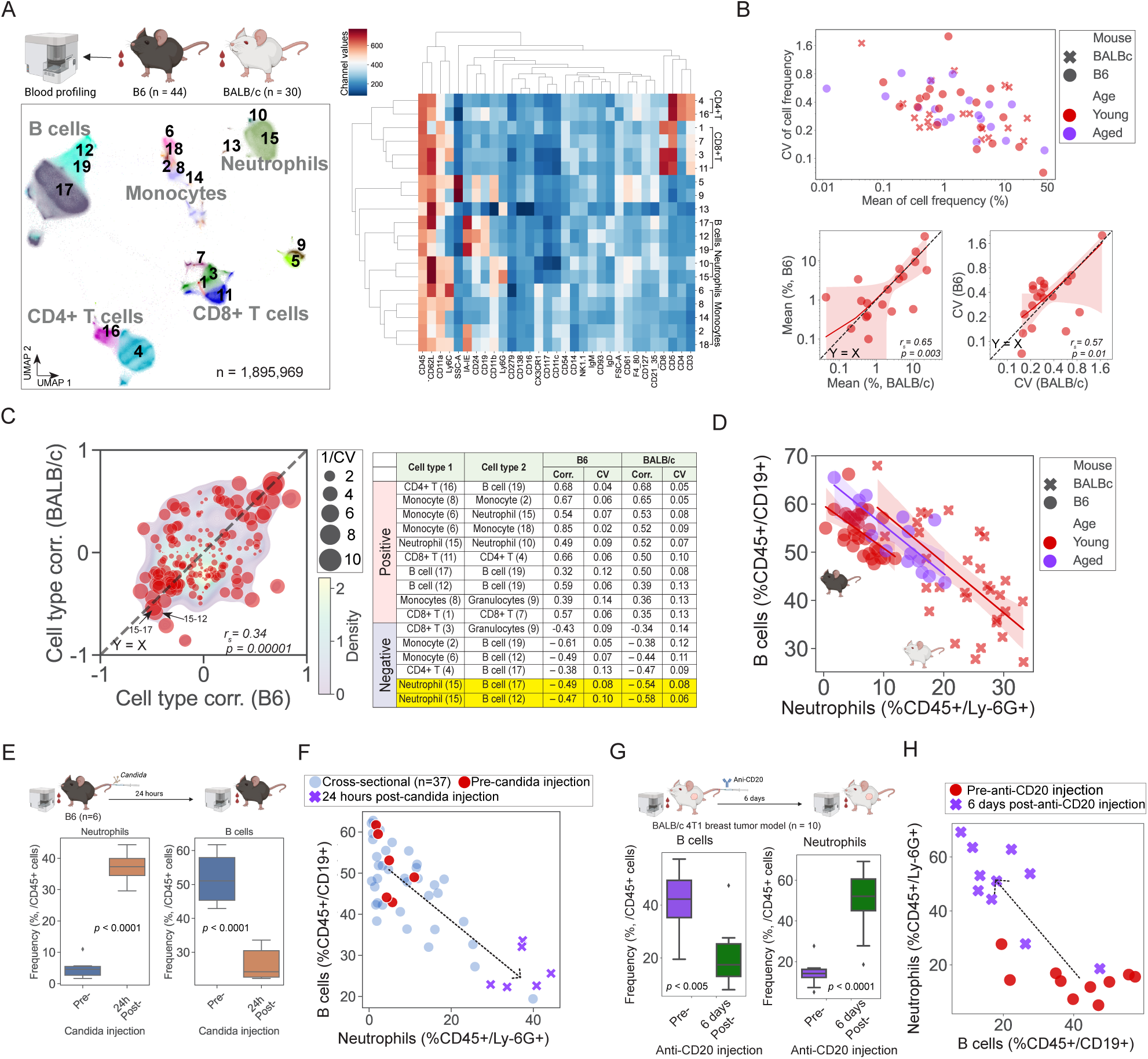
Blood profiling of B6 mice (n = 44) together with BALB/c (n = 30) mice demonstrates consistency of cell frequency variation and correlation. One to two drops of blood were acquired from 44 B6 and 30 BALB/c mice, followed by immediate processing/measurement by spectral flow cytometry. **(A)** UMAP visualization showing main blood cell types - neutrophils, B cells, monocytes, CD4+ and CD8+ T cells etc.(*left* ), and the hierarchical clustering result of marker expression showing 19 distinct cell types (*right* ). **(B)** Large fluctuations in cell frequencies is common across B6 (young and aged) and BALB/c mice (*top*). Both the mean frequency of the 19 cell types (*bottom left* ) and the variation (represented by CV) (*bottom right* ) are significantly positively correlated across B6 and BALB/c. The Spearman correlation result (*r_s_*) is shown in the corresponding figures. Each dot represents a cell type. Young mice were 6-week-old (red) Old mice were 26-week-old (blue). **(C)** Bootstrapping analysis of the cell frequency correlation (*left panel* ) The correlation coefficients for the frequencies of cells are themselves highly correlated when comparing B6 and BALB/c mice. Each dot represents the mean correlation coefficient (Corr.) of two cell types from 100 bootstrapping analysis. The size of the dot is proportional to the inverse of the sum of CV(Corr.) in B6 and BALB/c: the larger the dot, the less dispersed the mean is. The background is the kernel density estimate of all correlation coefficients between all different cell types from the bootstrapping analysis. (*right panel* ) Table summarizing few examples of pairs of cell types that exhibit strong correlations/anti-correlation with relatively low CVs in both B6 and BALB/c. **(D)** The strong anti-correlation between B cells and neutrophils is consistent across young B6 mice (n = 30, Spearman’s: *r_s_* = −0.56, *p <* 0.005; regression slope (95%), (−1.44, −0.34), old B6 mice (n = 14, Spearman’s: *r_s_* = −0.76, *p <* 0.005; regression slope (95%), (−1.49, −0.52) and young BALB/c mice (n = 30, Spearman’s: *r_s_* = −0.62, *p <* 0.001; regression slope (95%): (−1.50, −0.58). Each dot represents a mouse. (E) *Candida* infection in B6 mice leads to significant increase of blood neutrophils and concurrently decrease of blood B cells in 24 hours. **(F)** Anti-correlation of B cell and neutrophil frequency upon *candida* infection (*r_s_* = −0.81, *p <* 0.005). Linear regression analysis elucidates comparable slope (95%) (−1.02, −0.49) relative to the regression slope (95%) (−1.07, −0.55) identified by analyzing the CyTOF data of 39 healthy B6 mice (Fig. 1). The arrow connects the centroids (median of neutrophil and B cell frequency in each group) of the control and treatment groups.**(G)** Depletion of B cells in BALB/c 4T1 breast tumor mouse model by intraperitoneal injection of anti-CD20 leads to significant reduction of B cells and concurrently increase of neutrophils in the blood in 6 days. **(H)** Anti-correlation of B cell and neutrophil frequency upon B cell depletion (*r_s_* = −0.75, *p <* 0.001). Linear regression analysis of the B cell, neutrophils frequencies elucidates comparable slope (95%) (−1.48, −0.68) relative to the regression slope identified in the 30 healthy BALB/c mice in **(D)**. The arrow connects the centroids (median of neutrophil and B cell frequency in each group) of the control and treatment groups.

We next investigated the cell-cell frequency correlation that persisted across B6 (young, n=30) and BALB/c mice (young, n =30). We performed bootstrapping analysis of the cell type frequency correlation by sampling 28 mice from all 30 without replacement for 100 times (**Fig. 6C**). We found that the overall cell type correlation between B6 and BALB/c exhibited significant positive correlation (*r_s_*= 0.34, *p <* 0.00001), especially for the strong positively/negatively correlated cell pairs which almost align on the diagonal (**Fig. 6C**). Among the cell types that exhibited significant negative correlations, neutrophils and B cells were one of the most pronounced pairs in both B6 and BALB/c (**Figs. 6C-D**). The anti-correlation between neutrophils and B cells was so strong in both B6 and BALB/c that we could recapitulate their anti-correlation by just hand-gating the CD45^+^/CD19^+^ cells (B cells) and the CD45^+^/Ly-6G^+^ cells (neutrophils) (**Fig. 6D**). Relative to young B6 mice, aged B6 mice exhibited significantly higher mean of neutrophil frequency (two-sided *t* -test, *p <* 0.0001). Relative to B6 mice, BALB/c mice exhibited significantly higher mean of neutrophil frequency (*t* -test, *p <* 0.0001) and lower mean of B cell frequency (two-sided *t* -test, *p <* 0.0001). However, the anti-correlation between the neutrophils and B cells persisted in both B6 (young and old) and BALB/c.

Next we demonstrated that the robust anti-correlation between B cell and neutrophils predicted and accounted for the general opposite changes of B cells and neutrophils upon various perturbation/treatments in both healthy mice and mouse tumor models. We illustrate this idea in two experimental settings. In the first experiment, we investigated the response of neutrophils and B cells in B6 mice in response to an infection with *Candida albicans*. At 24 hours post-*candida* injection, we observed an average eight-fold increase of blood neutrophil frequency, consistent with prior studies and in agreement with the critical role of neutrophils in host defense against candidiasis ^30,31^ and a concurrent two-fold decrease of blood B cell frequency (**Fig. 6E**). The decrease of B cell frequency is not just a number game following the increase of neutrophil counts, as other cell types, such as blood macrophages (Ly6G*^−^*F4/80^+^) and CD4^+^ T cells (CD3^+^CD4^+^), exhibits significantly increased or decreased frequency to varying extents post-*candida* infection (**Fig. S29**). Significant anti-correlation between B cells and neutrophils was observed (*r_s_* = −0.81, *p <* 0.005) (**Fig. 6F**). The slope (95%) by linear regression analysis of B cell and neutrophil frequencies (−1.02, −0.49) is largely overlapping with the regression slope (95%) (−1.07, −0.55) identified by analyzing the CyTOF data of 39 healthy B6 mice (**Fig. 1**). In the second experiment, we directly depleted B cells in BALB/c 4T1 breast tumor mouse models by intraperitoneal injection of anti-CD20 antibody (**Fig. 6G**). We observed a significant reduction of blood B cell frequency, as expected, but also a concurrent increase of blood neutrophils at day 7 in the treatment group relative to control. Again, significant anti-correlation between B cells and neutrophils was observed upon treatment (*r_s_*= −0.75, *p <* 0.001). The 95% confidence interval on the slope (−1.48, −0.68) by regression analysis (**Fig. 6H**) is largely overlapping with the slope (95%) identified by analyzing 30 healthy BALB/c mice in (**Fig. 6D**) (−1.50, −0.58). The largely overlapping regression slopes (95%) in perturbation and in homeostasis uphold the promise to predict change in B cell numbers upon perturbing neutrophils and *vice versa*.

## DISCUSSION

The abundance of various cell types has been emerging prognostic markers for cancer outcomes and predictors of treatment response. For example, a recent study revealed that the B cell abundance, measured in either tumor tissues or peripheral blood mononuclear cells (PBMCs), is among the strongest predictors of immune checkpoint blockade (ICB) response in head and neck cancer patients ^32^. However, a quantitative understanding of the variability in cell numbers during hematopoiesis remains elusive, limiting the ability to optimize therapeutic interventions. Spontaneous oscillations of blood cell counts were first observed in leukemia and neutropenic patients about 50 years ago. The period of WBC oscillations varies in different patients with about 72 days in a 12-year old female CML patient^8^ and about 36 days in a 58-year old male CML patient^28^. The period of neutrophils oscillations in neutropenic patients varies from 18 to 24 days^29^. In addition to leukocytes, there is a broad involvement of other peripheral blood cell types including platelets, reticulocytes, erythrocytes etc., that exhibit fluctuating counts^7^. Not only in patients, the fluctuating leukocyte numbers have also been reported in healthy adults ^6,33,34^, which was mainly attributed to non-heritable influences through studies of twins at various ages ^34^. Despite recent progress to quantify hematopoiesis, measurements of cell frequency and its temporal dynamics are still missing ^35^. In this work, we systematically probed the dynamic features of hematopoiesis in healthy mice (B6 and BALB/c), by characterizing the variation, correlation of cell frequency and proliferation rates, and elucidating the potential chaotic nature of native hematopoiesis at homeostasis.

We showed that all cell types exhibit large variation of frequency in bone marrow, blood and spleen, and the variation of cell frequencies persisted across healthy B6 and BALB/c mice. The variation in cell frequencies contains rich information that can be used to deconvolve hematopoiesis Probabilistic frameworks have been developed to capture the variability and identify the most probable developmental path, exemplified by the identification of the development pathway of T cells: naive → central memory precursor → effector memory precursor → effector cells^15^. We observed that the variations in cell frequencies were not structured according to the hierarchical tree nature of hematopoiesis (**Fig. S8E**), suggesting that critical dynamic links were missing in our current understanding. Cell proliferation is one of the nonlinear processes that potentially would explain the large fluctuations of cell numbers. One striking observation from our study is that cell proliferation rates are indeed fluctuating-by-burst, and strongly correlated for all leukocytes in the bone marrow, blood and spleen.

We identified extensive positive and negative frequency correlations between various cell types in bone marrow, blood and spleen. One pronounced example is the strongly anti-correlated blood neutrophils and B cells in both B6 and BALB/c mice. The functional consequence is we now can predict how B cells and neutrophils respond to single perturbations. We showed that either direct B cell depletion in BALB/c mice or *candida* infection in B6 mice produces consistently anti-correlated changes of B cell and neutrophils, and the slope of their changes can be rationalized based on their anti-correlation identified in homeostasis. Such mutual inhibition between B cells and neutrophils can be inferred in other biological settings: reduction in B220^+^ cells in RAG1-deficient mice is compensated by increase of Gr-1^+^ cells but not by other cell types^36^. Circulating B cells can physically bind via CD18 to neutrophils in the lung microvasculature, and consequently drive apoptosis of neutrophils^37^. The correlation results identified by our study provide a starting point toward mechanistic understanding of cell-cell interaction and can be integrated during the design of lineage tracing toward a better understanding of the structure of hematopoiesis. It seems that no cell type is isolated and therefore targeting one cell type often leads to far-reaching changes for other cell types. This calls to caution when exploring immune therapies designed to target specific cell types.

With the continuous advancement of barcoding-based lineage tracing technology, a more nuanced picture of native hematopoiesis is being delineated. For example, the transposon barcoding study suggested that, in addition to the traditional lineage development, HSCs can directly generate megakaryocytes ^21^. The specific lineage commitments, *e.g.* divergence between erythrocytes and myeloid cells, have already started within the MPPs^38^. And it is the MPP but not the HSC that is responsible for long-term hematopoiesis^3^. Consistently, our experiments suggest MPPs exhibit significantly higher proliferation rates relative to HSCs on average (**Fig. S9D**). However, one limitation of the current barcoding-based lineage tracing study is that the textbook view of the hierarchical tree structure with fixed transition rates is still often used as the basis to design experiments and clinical trials, and this may blur our understanding of the interactions between differentiated cell types.

Cytokine-mediated cell-cell interaction is a critical feature of dynamic yet robust cell circuit design^24^. For example, the macrophages and fibroblasts can form a stable cell circuit through a “spring-and ceiling” mechanism^39^; the regulatory and effector T cells compete for cytokines to decide between tolerance and activation^40^ etc. Here we showed that a cytokine-mediated cell-cell interaction model is sufficient to generate deterministic yet chaotic behaviors. Such chaotic dynamics potentially explain both the large fluctuations and correlations for all cell types. Perturbing the cell proliferation rates, with the resulting increase/decrease of a particular cell type (B cells, myeloid cells, neutrophils etc.) in leukemia and neutropenic patients, can explain why the hematopoietic system transition from chaos to oscillations. More generally, chaotic dynamics have been observed in other biological systems, such as heart rate variability and neural activity, and conjectured to be the hallmark of reliability and resiliency in physiologically-healthy systems^25^. The loss of the chaotic behavior is indeed generally associated with pathological conditions. All together, our results support that hematopoiesis at homeostasis establish ‘information-rich’ chaotic dynamics at homeostasis. In contrast, highly periodic behaviors represent a loss of physiological complexity information and thus becoming fragile to perturbations. Our study uncovered fundamental properties of hematopoiesis, that may be critical for understanding the physiological variations of blood cell counts, for better diagnosing and monitoring various diseases, (such as anemia, leukemia, infections, and immune disorders), and for better designing perturbations in the context of immunotherapies.

### Limitations of the study

In the present study, we focused on characterizing the dynamics of hematopoiesis in two common inbred strains of laboratory mice (B6 and BALB/c). Future longitudinal studies of the hematopoietic contents in rewilded healthy mice and in healthy volunteers would provide further insights into how immune variation shapes responses to natural environment.

## Supplemental information index

There is one supplemental PDF, which includes Supplementary Sections 1-3, Figures S1-S28, Table S1-S3 and their legends in a PDF

## Acknowledgments

This work was supported by the intramural research program of the National Cancer Institute and in part by the division of intramural research of National Institute of Allergy and Infectious Diseases

## Author contributions

Conceptualization, D.J., R.M. and G.A-B.; methodology, D.J.,E. S-C, T.W, S-H.L., C.R., M. C-S., R.M and G.A.-B.; investigation, D.J.,E. S-C, T.W, S-H.L., C.R., M. C-S., R.M and G.A.-B; writing – original draft, D.J. and G.A-B.; writing – review & editing, all authors; funding acquisition, M.L., G.T. and G.A-B.; supervision, R.M. and G.A-B..

## STAR METHODS

### Key resources table Resource availability

#### Lead contact

Requests for further information and resources should be directed to and will be fulfilled by the lead contact, Grégoire Altan-Bonnet.

### Materials availability

#### Data and code availability

- All data reported in this paper will be shared by the lead contact upon request.
- All original codes of the mathematical model have been deposited at Zenodo ^41^ and is publicly available as of the date of publication.
- Any additional information required to reanalyze the data reported in this paper is available from the lead contact upon request.

